# Genome-wide identification of CDC34 that stabilizes EGFR and promotes lung carcinogenesis

**DOI:** 10.1101/255844

**Authors:** Xin-Chun Zhao, Gui-Zhen Wang, Yong-Chun Zhou, Liang Ma, Jie Liu, Chen Zhang, Da-Lin Zhang, San-Hui Gao, Li-Wei Qu, Bin Zhang, Chang-Li Wang, Yun-Chao Huang, Liang Chen, Guang-Biao Zhou

## Abstract

To systematically identify ubiquitin pathway genes that are critical to lung carcinogenesis, we used a genome-wide silencing method in this study to knockdown 696 genes in non-small cell lung cancer (NSCLC) cells. We identified 31 candidates that were required for cell proliferation in two NSCLC lines, among which the E2 ubiquitin conjugase CDC34 represented the most significant one. CDC34 was elevated in tumor tissues in 67 of 102 (65.7%) NSCLCs, and smokers had higher CDC34 than nonsmokers. The expression of CDC34 was inversely associated with overall survival of the patients. Forced expression of CDC34 promoted, whereas knockdown of CDC34 inhibited lung cancer *in vitro* and *in vivo*. CDC34 bound EGFR and competed with E3 ligase c-Cbl to inhibit the polyubiquitination and subsequent degradation of EGFR. In EGFR-L858R and EGFR-T790M/Del(exon 19)-driven lung cancer in mice, knockdown of CDC34 by lentivirus mediated transfection of short hairpin RNA significantly inhibited tumor formation. These results demonstrate that an E2 enzyme is capable of competing with E3 ligase to inhibit ubiquitination and subsequent degradation of oncoprotein substrate, and CDC34 represents an attractive therapeutic target for NSCLCs with or without drug-resistant EGFR mutations.

## Introduction

The ubiquitin (Ub)-proteasome system (UPS) is the principal pathway for diverse intracellular protein degradation, in which the E2 ubiquitin-conjugating enzymes play critical roles by transferring the Ub on their conserved cysteine residue to the ε-amino group of lysine residues on substrates^1^. The cell division cycle 34 (CDC34, or UBCH3, Ubc3, UBE2R1) is an E2 enzyme which contains a non-covalent Ub binding domain in its carboxyl terminus^2^ and employs distinct sites to coordinate attachment of Ub to a substrate and assembly of polyubiquitin chains^3^. CDC34 functions in conjunction with the Skp1-Cullin 1-F-box (SCF) E3 Ub ligase to catalyze covalent attachment of polyubiquitin chains to substrates such as p27^4^, ATF5^5^, WEE1^6^, IκBα^7^, Grr1^8^, c-Ski^9^, Sic1^10^, Far1^11^, and other substrate proteins. However, whether CDC34 and other E2s have biological functions besides E2 activity remains to be investigated.

Hyperactivation of the epidermal growth factor receptor (EGFR) by gain-of-function mutations and overexpression have been found in more than a half of patients with non-small cell lung cancers (NSCLCs)^12, 13^, and inhibition of constitutively activated EGFR significantly benefits lung adenocarcinoma patients with mutant EGFR^14^. The proteolysis of EGFR protein is controlled by an E3 ligase c-Cbl and E2 conjugase UbcH7 ^15^ or Ubc4/5^16^. c-Cb1 mediates the ubiquitination, endosome fusion, and lysomal sorting of EGFR^17, 18, 19^. The conserved N-terminal of c-Cbl is sufficient to enhance EGFR ubiquitination^20^, while the RING finger C-terminal flank controls EGFR fate downstream of receptor ubiquitination^21^. Somatic mutations and loss of heterozygosity (LOH) of c-Cbl had been reported in a proportion of NSCLCs^22^, how EGFR evades proteolytic degradation and thus accumulates in lung cancer remains to be elucidated.

Abnormalities in ubiquitin pathway genes (UPGs) have been reported in lung cancer. These alterations include E1 ubiquitin activating emzyme UBE1L^23^, E2 ubiquitin-conjugating enzymes UbcH10^24^, UBE2C^25^, Hrad6B^26^, E3 ligases^27^ such as c-Cbl^22^, SINA^28^, deubiquitylases Ataxin-3^29^, USP8^30^, USP17^31^, USP37^32^, and SCF essential component Skp1^33^. To systematically identify UPGs that are crucial for lung cancer cells proliferation and growth, we conducted a genome-wide silencing of the E1, E2, E3, and deubiquitinases in NSCLC cell lines. We reported that 31 UPGs were required for the survival of two NSCLC lines, among them CDC34 was overexpressed and inversely associated with clinical outcome of NSCLC patients. Interestingly, CDC34 competitively bound EGFR and prevented it from proteolysis to promote lung cancer.

## Results

### A genome-wide silencing of UPGs in lung cancer cells

A total of 696 UPGs (Supplementary Table 1) were silenced by transfection of small interfering RNA (siRNA) of the Dharmacon human siGENOME SMARTpool library into the NSCLC lines A549 (with wild type EGFR) and H1975 (harboring L858R/T790M EGFR mutation). Cell viability was measured 72 h after transfection, and Z-scores from duplicate experiments for each SMARTpool (Fig. 1a) were determined^34^. To validate the results, a secondary screen was performed in A549 and the results showed that the robust Z scores of most of the genes across repeats were strongly correlated (r=0.86; Fig. 1b). A SMARTpool was considered a hit if the Z-score was ≤ -2 both in A549 and H1975 (Fig. 1a and Supplementary Table 1). Eighty hits (11.5%) were identified in A549 and 72 hits (10.3%) were uncovered in H1975 cells (Supplementary Fig. 1a), with 31 hits discovered in both cells (Supplementary Fig. 1a and Fig. 1c). These hits include two E2 enzymes (UBE2T, CDC34), twenty E3 ligases (UBE3A, WWP2, TRIM68, and others), and nine others (e.g. USP48, PSMD14, OUTD5).

**Fig. 1.**
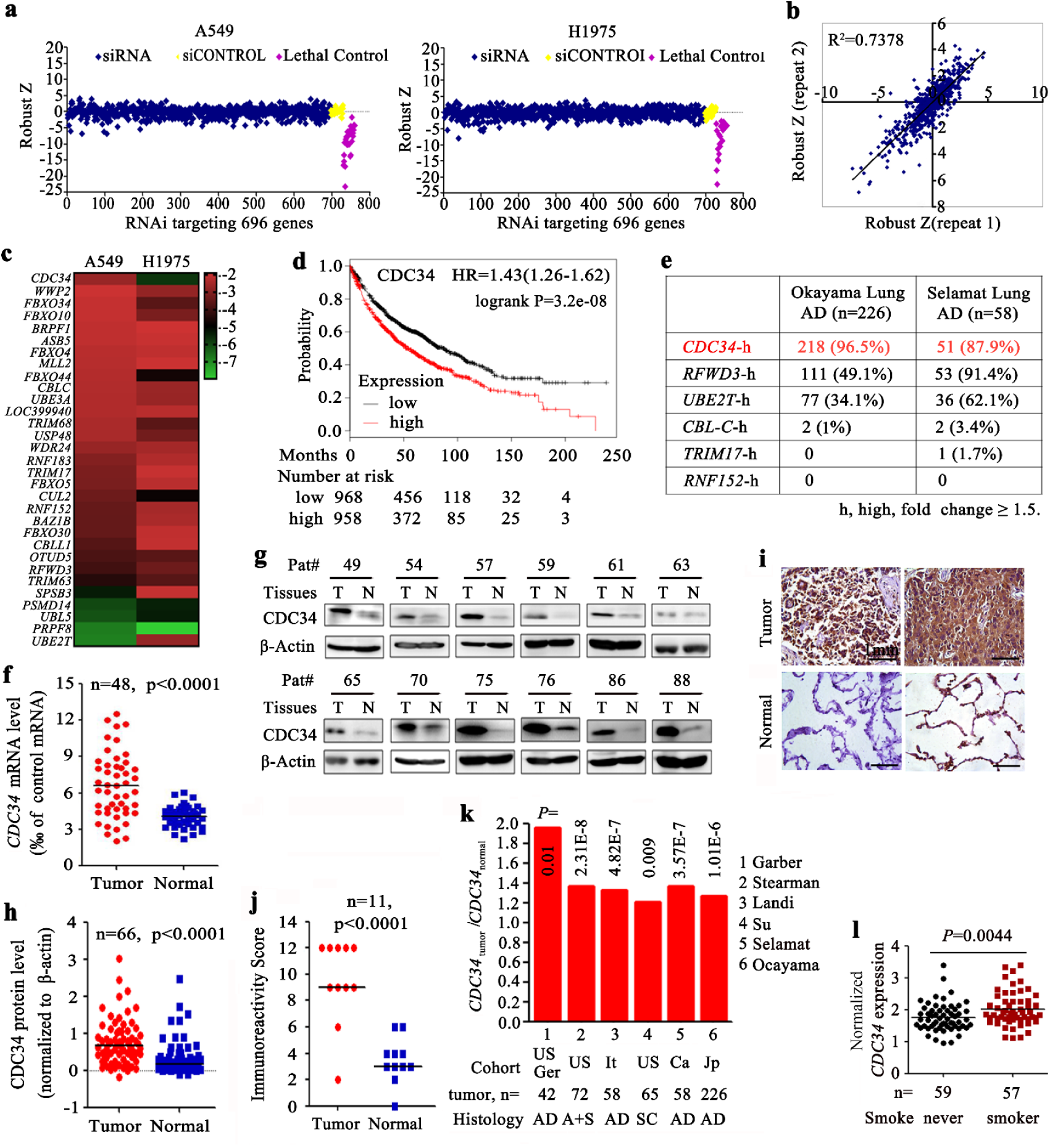
Identification of *CDC34* as a crucial oncogene in NSCLCs by siGenome screening. (a) A549 and H1975 cells were treated with 50 nM siRNAs from the siGenome library (containing 696 UPGs) for 72 hours and the cell viability was measured. The z score was calculated as described in materials and methods. (b) A second replicated screening was conducted in A549 to validate the results of the initial screening. (c) Heat map showing 31 candidates with Z score ≤ −2 in both A549 and H1975. (d) Overall survival of NSCLCs with high or low expression of *CDC34*. (e) The expression of 6 candidate genes in Okayama and Selamat cohorts (Tumor/Normal ≥ 1.5). (F-K) The expression of *CDC34* was tested by qRT-PCR (f), Western blot (g, h), and immunohistochemistry (i, j) assays in NSCLCs of our cohort. Size bar, 1 mm. The densitometry analysis of the Western blot results (h) and the immunoreactivity score (j) was calculated. (k) *CDC34* expression was detected by microarrays in tumor samples and normal lung tissues in Oncomine datasets. AD, adenocarcinoma; SC, squamous cell carcinoma; A+S, adenocarcinoma and squamous cell carcinoma; Ger, Germany; Ca, Canada; It, Italy; Jp, Japan; Tw, Taiwan, China. (l) *CDC34* in never smoker and smoker NSCLCs of the work of Selamat et al ^37^ in Oncomine datasets.

To identify genes that are critical to lung carcinogenesis, the association between the expression of the 31 genes and the clinical outcome of NSCLC patients was analyzed using the Online Survival Analysis Software^35^ (<u>http://kmplot.com/analysis/index.php?p=service&cancer=lung</u>). We reported that the expression of six genes, *CDC34* (Fig. 1d), *UBE2T*, *RNF152*, *TRIM17*, *CBLC*, and *RFWD3*, was inversely associated with overall survival of the patients. The data^36, 37^ of the cancer microarray database Oncomine^38^ (<u>http://www.oncomine.org</u>) showed that, of these 6 genes, *CDC34* was most significantly upregulated in lung tumors compared to normal tissues (Fig. 1e). CDC34 was therefore chosen for further study.

### Overexpression of CDC34 in NSCLCs

We tested the expression of CDC34 in 102 NSCLCs (Table 1) by quantitative reverse transcription polymerase chain reaction (qRT-PCR), and showed that in 67 (65.7%) of the patients the expression of CDC34 was significantly elevated in tumor samples than the counterpart normal lung tissues (Fig. 1f). A two-sided Fisher exact test showed that the expression of CDC34 in smoker NSCLCs was significantly higher than in non-smoker patients (*P*=0.027; Table 1). Western blot (Fig. 1g, h) and immunohistochemistry (IHC) (Fig. 1i, j) assays showed the elevation of CDC34 in tumor samples. In works of Oncomine database^37, 39, 40, 41, 42, 43^, *CDC34* expression in tumor samples was higher than in their paired normal lung tissues (Fig. 1k), and smokers^37^ had higher *CDC34* expression in tumor tissues than nonsmokers (Fig. 1l).

**Table 1.** Summary of baseline demographic characteristics of the 102 patients.

| **Characteristics** | **Cases,n** | **CDC34 high, n (%)** | ***P* values\*** |
| --- | --- | --- | --- |
| **Total number** | **102** | **67 (65.7)** |  |
| **Age** |  |  |  |
| <65 | 74 | 51 (68.9) | 0.264 |
| ≥65 | 28 | 16 (57.1) |  |
| **Gender** |  |  |  |
| male | 71 | 50 (70.4) | 0.127 |
| female | 31 | 17 (54.8) |  |
| **Smoking** |  |  |  |
| smoker | 59 | 44 (74.6) | 0.027 |
| non-smoker | 43 | 23 (53.5) |  |
| **Histology** |  |  |  |
| adenocarcinoma | 61 | 36 (59) | 0.093 |
| squamous-cell carcinoma | 37 | 28 (75.7) |  |
| others | 4 | 3 (75) |  |
| **TNM stage** |  |  |  |
| I-II | 58 | 35 (60.3) | 0.219 |
| III-IV | 43 | 31 (72.1) |  |
| no record | 1 | 1 (100) |  |

\*, *P* values were calculated using a two-sided Fisher's exact test.

### CDC34 is required for lung cancer cells proliferation

To evaluate the role of *CDC34* in lung cancer, siRNA against *CDC34* (si*CDC34*) was transfected into A549 and H1975 cells, which resulted in a 50-90% reduction of CDC34 protein (Fig. 2a) in the two lines. We found that silencing of CDC34 led to a significant inhibition of cell growth/proliferation (Fig. 2a, b) and suppression of colony forming activity (Fig. 2c) of the cells. On the contrary, exogenous expression of CDC34 significantly increased the proliferation (Fig. 2d), growth (Fig. 2e), and colony forming activity (Fig. 2f) of the cells.

**Fig. 2.**
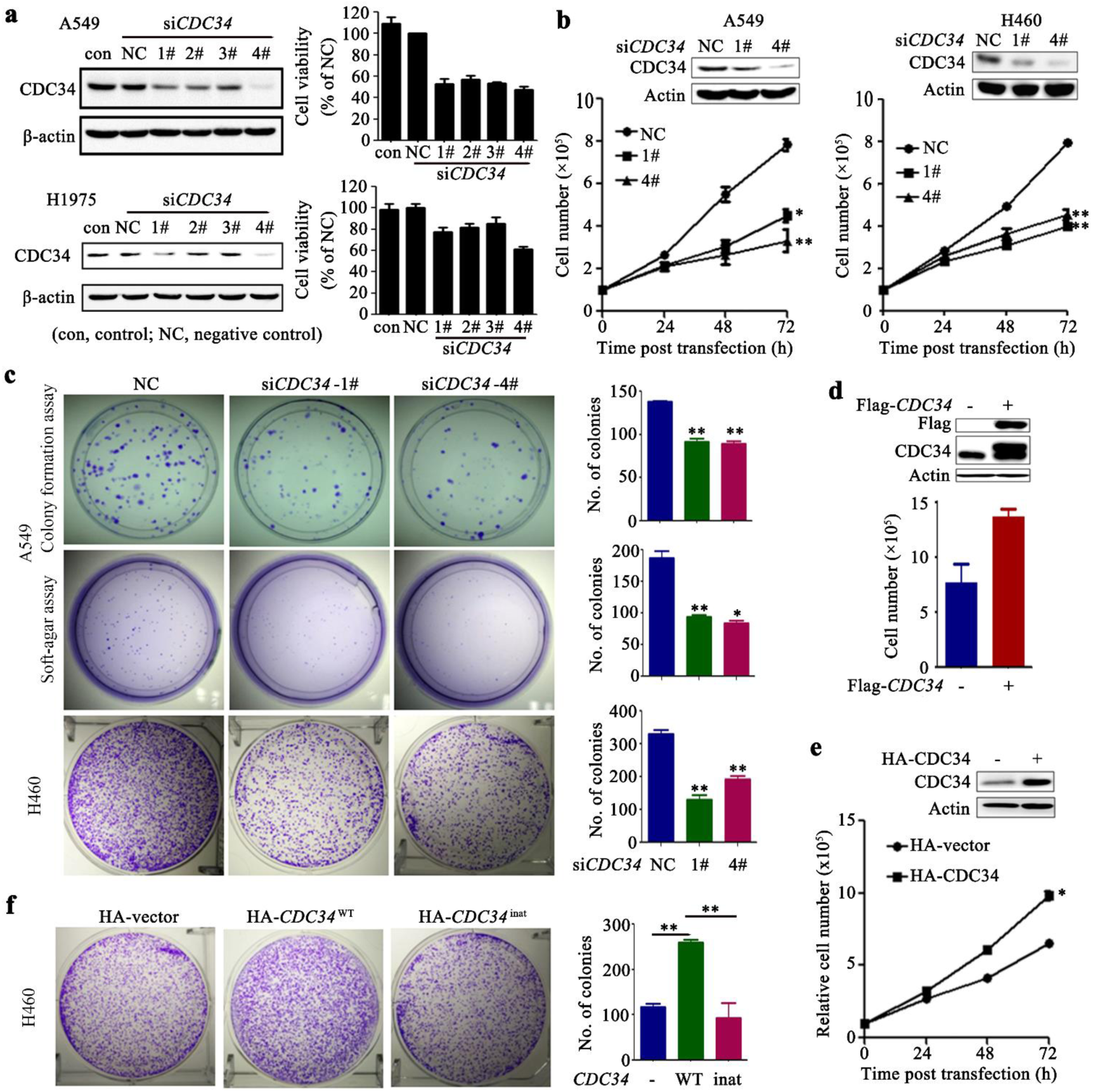
CDC34 is required for proliferation of NSCLC cells *in vitro* and *in vivo*. (a) A549 and H1975 cells were transfected with si*CDC34* (1# to 4#) and cell viability was assessed by the CellTiter-Glo Reagent. The relative expression of CDC34was detected by immunobloting 72 h after si*CDC34* transfection. (b) A549 and H460 cells were transfected with si*CDC34* (1# and 4#) and cell proliferation was assessed by trypan blue exclusion analysis. The relative expression of CDC34 was detected by Western blot. (c) Colony formation and soft-agar assays of A549 and H460 cells transfected with si*CDC34*. (d-f) The H460 cells were transfected with Flag or HA-tagged *CDC34* and cell proliferation was assessed by trypan blue exclusion analysis or colony formation assays. NC, negative control. The error bars indicate the SD of three independent experiments; \**P*< 0.05, \*\**P*< 0.01.

### CDC34 positively regulates EGFR

We aimed to identify the downstream target of CDC34 in lung cancer. By using the data of the reverse phase protein array of 235 lung adenocarcinomas from a previous work^44^, we analyzed the potential association between *CDC34* and some driver proteins of lung cancer such as EGFR, N-RAS, MEK1, and LKB1, as well as CDC34’s substrate protein p27 ^45^. We reported that the expression of *CDC34* was inversely associated with p27 expression (*P*=0.0127). Interestingly, CDC34 expression level was also correlated with EGFR (*P*=0.0015, Fig. 3a), suggesting that CDC34 may have a role in determining EGFR expression. IHC assays of NSCLC samples showed that patients with higher CDC34 had higher EGFR in the tumor tissues (Fig. 3b). We then showed that silencing of CDC34 in H460 and A549 cells resulted in downregulation of EGFR (Fig. 3c, Supplementary Fig. 2a). In H1975 and HCC827 (with an E746-A750 in-frame deletion of EGFR) cells, si*CDC34* also downregulated EGFR (Fig. 3c and Supplementary Fig. 2b). Moreover, knockdown of CDC34 resulted in inhibition of EGF-induced phosphylated EGFR (pEGFR) (Fig. 3c, Supplementary Fig. 2a) and constitutively activated EGFR (Fig. 3c and Supplementary Fig. 2b), and suppressed its kinase activity (Fig. 3d). Constistently, pERK1/2, pAKT and pSTAT3 were also dowregulated by si*CDC34* (Fig. 3e, Supplementary Fig. 2c). On the contrary, exogenous CDC34 expression significantly increased protein levels of EGFR/pEGFR (Fig. 3f, S2d) and kinase activity (Fig. 3g). Besides, the reduction of EGFR caused by silencing CDC34 was rescued by overexpressed CDC34 whose CDS region was synonymously mutated in case of targeting by siRNA against *CDC34* (CDC34^res^; Fig. 3h).

**Fig. 3.**
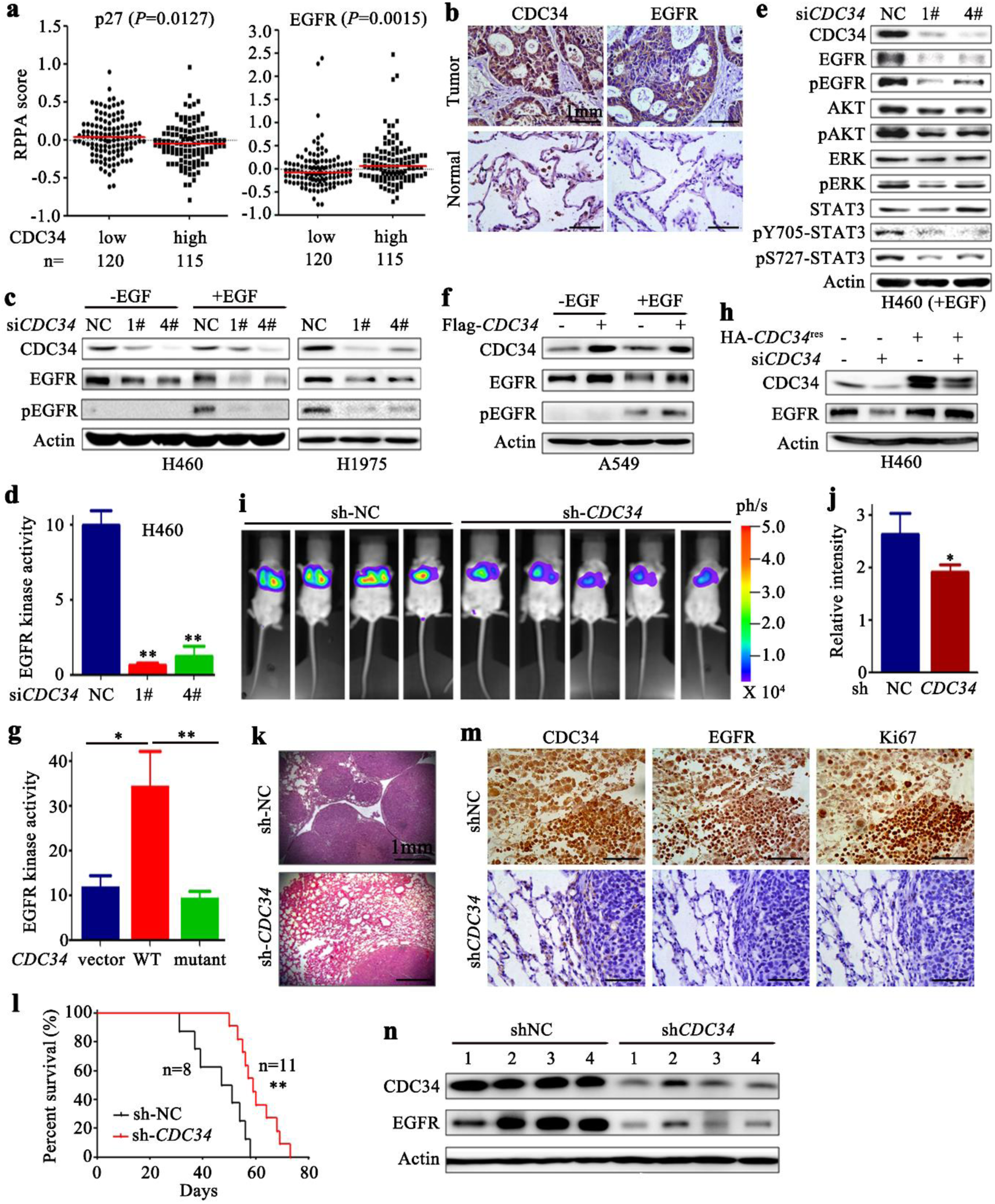
CDC34 positively regulates EGFR. (a) A scatter diagram of reverse phase protein array (RPPA) data showing a negative correlation between the mRNA level of *CDC34* and protein level of p27, and a positive correlation between the mRNA level of *CDC34* and protein level of EGFR. The data are from TCGA datasets. (b) IHC analysis of CDC34 and EGFR in NSCLCs of our setting. (c) Western blot analysis of EGFR/p-EGFR in cells transfected with si*CDC34* in H460 and H1975 cells with or without EGF co-incubation. (d) H460 cells transfected with si*CDC34* were lysed, the lysates were immunoprecipitated with an anti-EGFR antibody and assayed for tyrosine kinase activity. Error bars, standard deviation. (e) Western blot analysis of EGFR/pEGFR and downstream molecules in EGF-stimulated, si*CDC34*-transfected H460 cells. (f) Immunoblotting of lysates of Flag-*CDC34*-expressing A549 cells co-incubated with or without EGF. (g) A549 cells transfected with *CDC34* were lysed, the lysates were immunoprecipitated with an anti-EGFR antibody and assayed for tyrosine kinase activity. Error bars, standard deviation. (h) Immunoblot assay of EGFR in cells transfected with si*CDC34* in the absence or presence of *CDC34^res^*, which is mutated to resist si*CDC34* targeting. (i) SCID mice were injected with 1×10 ^6^ A549-luciferase cells via tail vein, and 30 days later the mice were detected by the IVIS Spectrum system. (j) The relative luciferase intensity in the mice. (k) Hematoxylin-eosin staining of the lung sections from mice of each group. (l) Kaplan– Meier survival curve of the mice. (m) IHC analysis of CDC34, EGFR and Ki67 in tumors from mice of each group. (n) The expression of CDC34 and EGFR in the tumor samples from the mice of each group.

### Knockdown of CDC34 suppresses lung cancer cells growth *in vivo*

To study the effect of CDC34 on lung cancer cells proliferation *in vivo*, A549-luciferase cells transfected with sh*CDC34* (Supplementary Fig. 2e) were inoculated into SCID-beige mice via tail vein, and the results showed that the luciferase signal in CDC34 knockdown groups was significantly lower than in the control group (Fig. 3i, j). Hematoxylin-eosin (HE) staing showed that the lungs from the control group were almost full of tumor cells, but lungs from the CDC34 knockdown mice had markedly less tumor cells (Fig. 3k). Moreover, the overall survival of the mice harboring the CDC34-knockdown cells was significantly longer than the control group (Fig. 3l). The expression level of CDC34 and EGFR was tested in the tumor tissues by IHC and Western blot assays, and the results showed that in the lungs of the mice inoculated with A549-luciferase-sh*CDC34* cells, the protein level of the two (Fig. 3m, n) was markedly lower than the control group. In addition, Ki67 staining in CDC34 knockdown cells was drastically decreased compared to tumor tissues of control group mice (Fig. 3m). These findings demonstrated that knockdown of CDC34 suppresses lung cancer cell growth *in vivo* by inhibiting EGFR.

### CDC34 interacts with EGFR

To elucidate the mechanism of CDC34 in regulating EGFR, we performed CO-immunoprecipitation (CO-IP) assay and found that EGFR was precipitated by a monoclonal anti-CDC34 antibody in both H460 and H1975 cells (Fig. 4a). Consistently, CDC34 was detected in a monoclonal anti-EGFR antibody-precipitated proteins (Fig. 4a). The immunofluorescence analysis showed that CDC34 colocalized with EGFR mainly in intracelluar region near the cell membrane (Fig. 4b). In 293T cells, ectopic expressed Flag-EGFR interacted with CDC34 (Fig. 4c). To confirm these findings, purified GST-CDC34 was co-incubated with lysates of H460 and H1975 cells, and precipitated EGFR was detected by immunoblot (Fig. 4d). On the other hand, the purified GST-EGFR also precipitated CDC34 from H460 cell lysates (Fig. 4e). In H460 and 293T cells, CDC34 was capable of interacting with EGFR in the presence and absence of EGF stimulation (Supplementary Fig. 3a, b), suggesting that CDC34-EGFR interaction was independent of the phosphorylation status of EGFR.

**Fig. 4.**
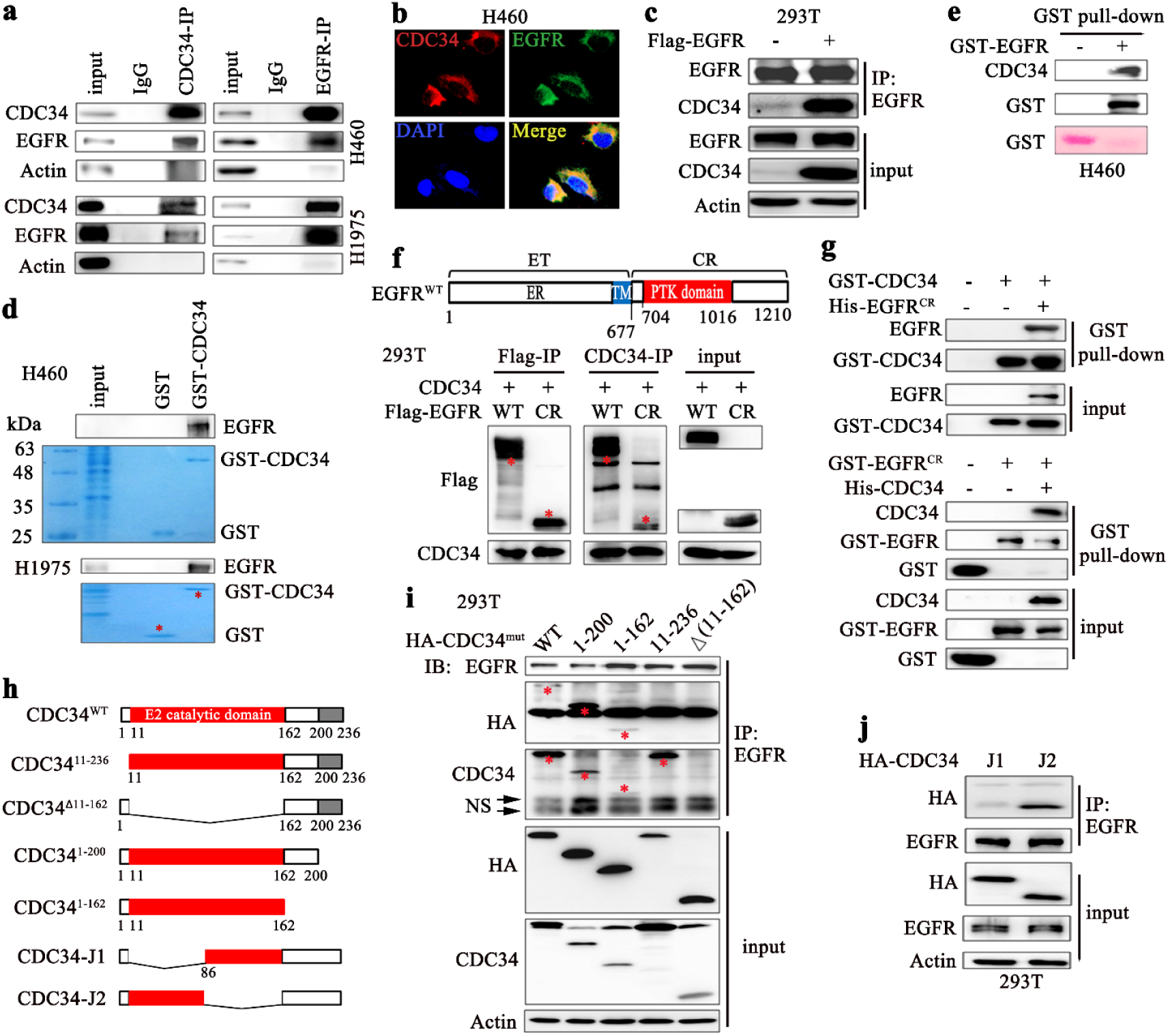
CDC34 interacts directly with EGFR. (a) Co-IP and immunoblotting assays using indicated antibodies and cell lysates. (b) Immunofluorescence assay of H460 cells using antibodies against CDC34 (red) and EGFR (green), and 4’,6-diamidino-2-phenylindole (DAPI) to counterstain the nucleus (blue). (c) Co-IP and immunoblotting assays using indicated antibodies and lysates of Flag-EGFR-transfected cells. (d) GST or GST-CDC34 was incubated with lysates of NSCLC cells *in vitro* to detect endogenous EGFR using indicated antibodies. (e) GST or GST-EGFR was incubated with lysates of H460 *in vitro* to detect endogenous CDC34 using indicated antibodies. (f) The cells were transfected with Flag-*EGFR*, lyzed, and Co-IP and immunoblot assays were performed using indicated antibodies. Schematic representation of EGFR proteinis shown at upper panel. (g) *In vitro* Co-IP and immunoblot experiments using purified CDC34 and EGFR proteins and indicated antibodies. GST‐ and His-tagged proteins were isolated from 293T cells that were transfected with respective plasmids. (h) Schematic representation of CDC34 truncted mutants. (i) Co-IP and immunoblotting assays using indicated antibodies and lysates of HA-*CDC34*-transfected cells. (j) Co-IP and immunoblotting assays using lysates of HA-*CDC34*-transfected cells and indicated antibodies.

To determine the CDC34-binding region within EGFR, several deletion mutants were constructed, and the results showed that the intracellular domain (EGFR^CR^) could bind CDC34 in cells (Fig. 4f). *In vitro* assays using purified GST-CDC34 and His-EGFR^CR^ (Fig. 4g, upper panel) or GST-EGFR^CR^/His-CDC34 (Fig. 4g, lower panel) confirmed the direct binding of the two proteins. To unveil the region of CDC34 that binds to EGFR, a series of deletion mutants were designed (Fig. 4h) and transfected into 293T cells. We found that only the mutant lacking E2 catalytic domain could not bind to EGFR (Fig. 4i). Further deletion assays showed that the first half of the E2 catalytic domain (K11 – I86) was the binding site of CDC34 for EGFR (Fig. 4j). However, neither the active site residue Cys93 nor the C-terminal acidic domain of CDC34 was required for interaction with EGFR, since mutations in these sites did not affect CDC34-EGFR binding affinity (Fig.S3c).

### CDC34 protects EGFR from proteolytic degradation

We noticed that knockdown of CDC34 in A549 and H1975 cells resulted in downregulation of EGFR at protein (Fig. 3c, e) but not mRNA level (Fig. 5a), suggestive of a role for CDC34 in stabilizing EGFR. The cycloheximide (CHX) chase assay showed that the half-life of EGFR decreased upon knockdown of CDC34 (Fig. 5b), but increased upon forced expression of CDC34 (Supplementary Fig. 4a).

**Fig. 5.**
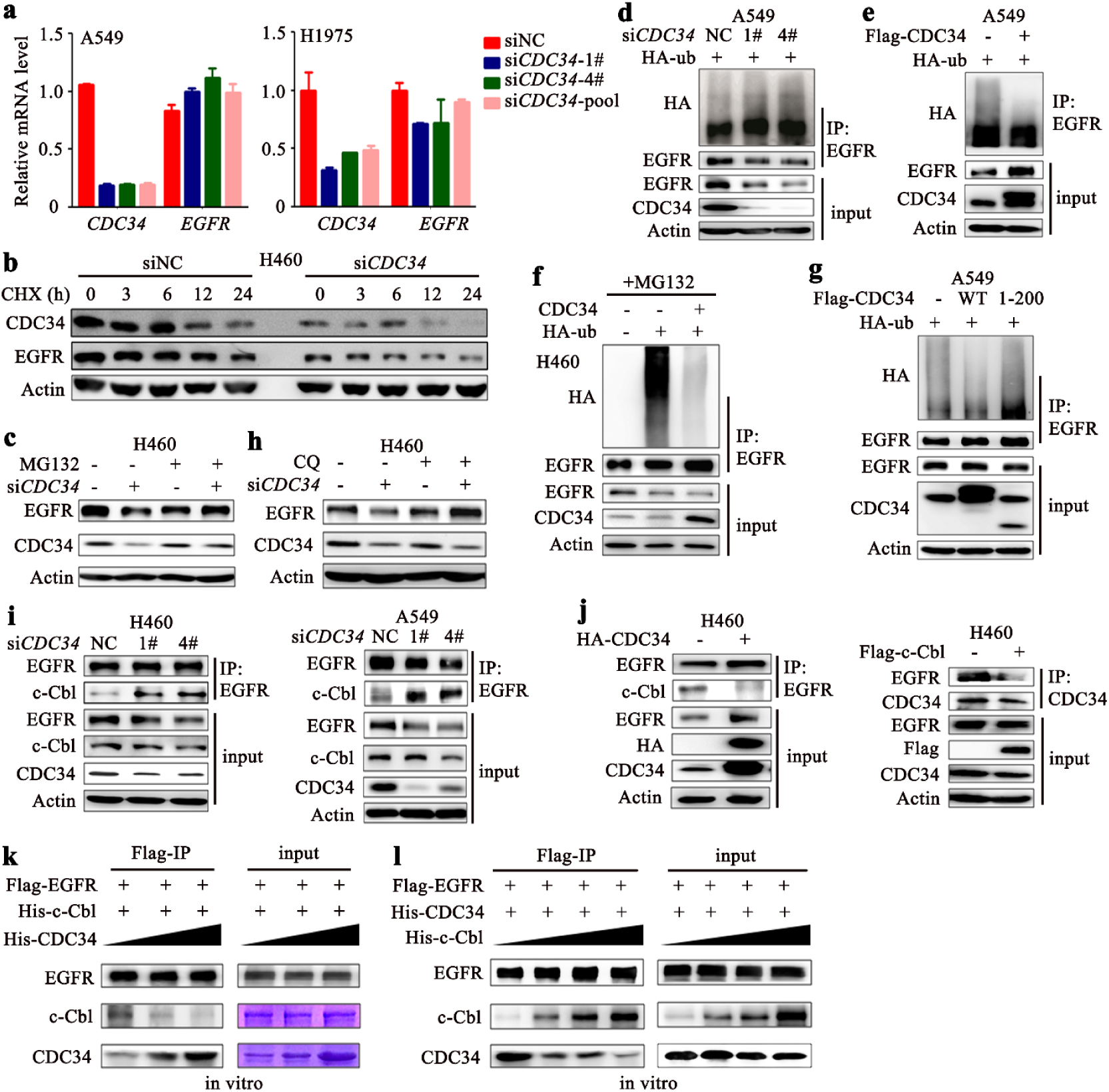
CDC34 inhibits the ubiquitination of EGFR and protects it from proteolytic degradation. (a) qRT-PCR analysis of *CDC34* and *EGFR* in A549 and H1975 cells 72 h after si*CDC34* transfection. (b) Cycloheximide (CHX) chase assay to measure EGFR half-life in H460 cells transfected with si*CDC34*. (c) Western blot analysis of the EGFR expression in H460 cells transfected with si*CDC34* in the presence or absence of MG132. (d) A549 cells were transfected with si*CDC34* and HA-Ub, lysed, and the lysates were subjected to Co-IP and immunoblot using indicated antibodies. (e) A549 cells were transfected with Flag-*CDC34* and HA-Ub, and lysed for Co-IP and immunoblot assays. (f) H460 cells were transfected with indicated constructs, treated with MG132, and lysed for Co-IP and immunoblot assays. (g) The cells were transfected with CDC34 and HA-Ub, and lysed for Co-IP and immunoblot assays using indicated antibodies. (h) Western blot analysis of EGFR in si*CDC34*-transfected H460 cells in the presence or absence of chloroquine (CQ). (i) H460 and A549 cells transfected with si*CDC34* were lysed and subjected to Co-IP and immunoblotting using indicated antibodies. (j) H460 cells were transfected with *CDC34* (left) or *c-Cbl* (right) for 48 h, and lysed for Co-IP and immunoblot using indicated antibodies. (k) *In vitro* Co-IP experiments using purified Flag-EGFR, His-c-Cbl, and increasing amount of purified His-CDC34 proteins and indicated antibodies. (l) *In vitro* Co-IP experiments using purified Flag-EGFR, His-CDC34, and increasing amount of purified His-c-Cbl proteins and indicated antibodies.

Previous studies showed that EGFR turnover was controlled by proteasome and lysosome pathways. We found that the proteasome inhibitors MG132 effectively inhibited si*CDC34*-induced EGFR down-regulation (Fig. 5c). Indeed, silencing of endogenous CDC34 increased the ubiquitination level of EGFR (Fig. 5d), whereas exogenous expression of CDC34 reduced its ubiquitination at the absence (Fig. 5e) or presence of MG132 (Fig. 5f). Furthermore, compared with wild-type CDC34, a mutant transcript CDC34^1-200^ failed to suppress EGFR ubiquitination in the absence (Fig. 5g) and presence of MG132 (Supplementary Fig. 4b). The lysosome inhibitor chloroquine (CQ) also inhibited si*CDC34*-caused downregulation of EGFR at protein level (Fig. 5h).

c-Cbl was reported to mediate the degradation of EGFR ^46^. By using the CO-IP experiment, we found that si*CDC34* drastically enhanced c-Cbl-EGFR interaction in H460 and A549 cells (Fig. 5i). In contrast, the protein level of c-Cbl that binds EGFR reduced upon CDC34 overexpression (Fig. 5j, left panel). Then we overexpressed c-Cbl in H460 cells, and observed the decreased EGFR-CDC34 binding affinity (Fig. 5j, right panel). In *in vitro* experiments using purified proteins, we found that increased levels of CDC34 supressed c-Cbl-EGFR binding affinity (Fig. 5k), while increased dosages of c-Cbl diminished CDC34-EGFR binding affinity (Fig. 5l, Supplementary Fig. 4c), indicating that CDC34 may compete with c-Cbl to bind EGFR and protects it from ubiquitination and subsequent degradation.

### Knockdown of CDC34 suppresses EGFR L858R-driven lung cancer

To evaluate the therapeutic potentials of CDC34 inhibition, an EGFR^L858R^-driven lung cancer mouse model was established as described^47^, and the lentiviral particles containing short hairpin RNA targeting *CDC34* (sh*CDC34*) were generated and intranasally administrated into the mice lungs before the mice were treated with doxycycline (DOX) to induce lung cancer ^47^. One month after the initiation of DOX treatment, the mice were detected by microscopic computed tomography (micro-CT). We found disseminated tumors in the lungs of control group mice, but only small tumors were seen in the *CDC34* knockdown group mice (Fig. 6a). Histological examination of the lungs demonstrated that the sh*CDC34*-treated mice developed lesions in alveoli, whereas the shNC group mice harbored disseminated adenocarcinomas (Fig. 6b). qRT-PCR, immunoblot, and IHC assays of the lung specimens revealed the downregulation of CDC34 at both mRNA and protein levels (Fig. 6b) and the decrease in EGFR as well as Ki67 (Fig. 6c) in sh*CDC34*-treated mice. Western blot analysis further showed that DOX administration induced the expression of EGFR, and silencing of CDC34 downregulated EGFR in lungs of the mice (Fig. 6d). In addition, the overall survival of the sh*CDC34* group mice was significantly prolonged as compared with shNC-treated mice (Fig. 6e).

**Fig. 6.**
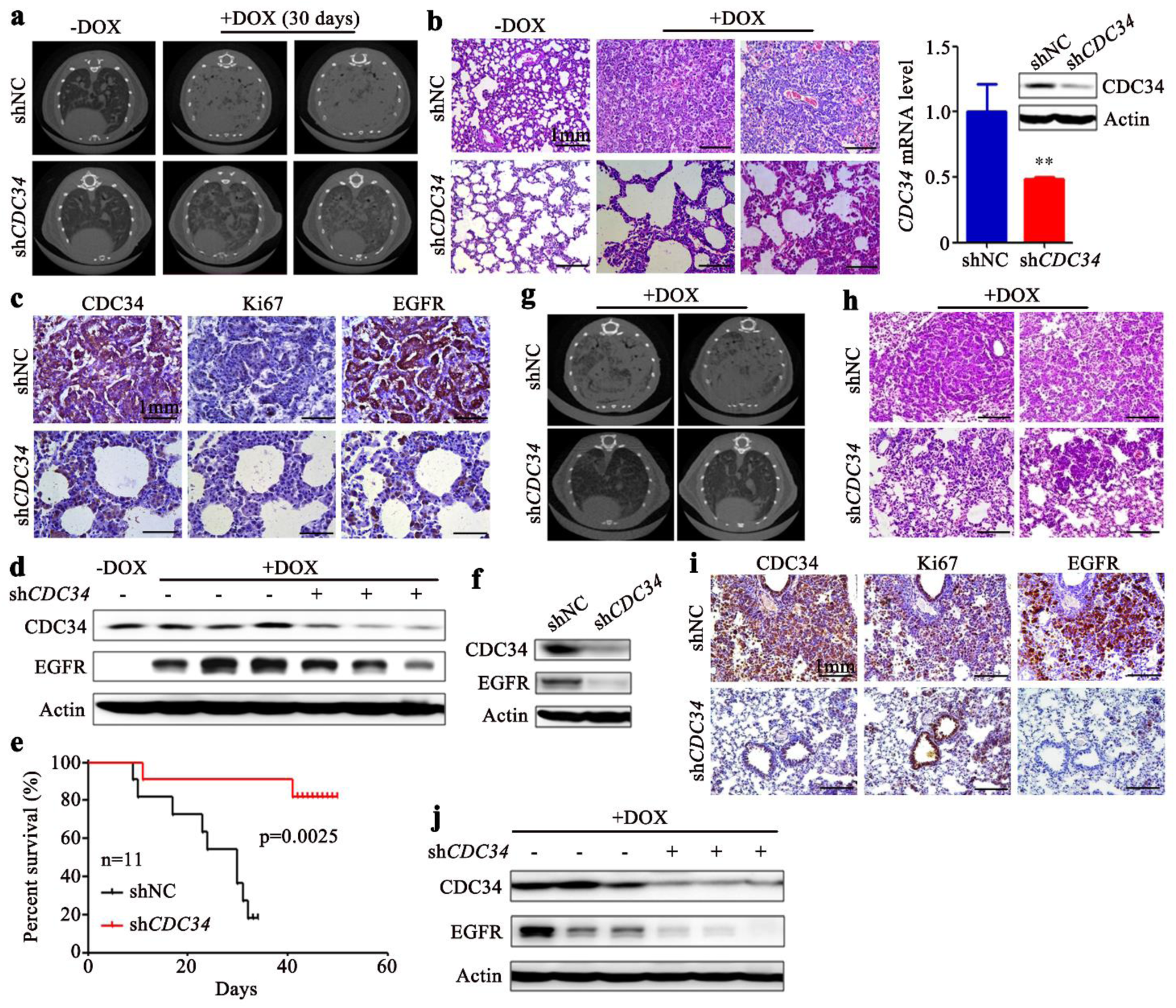
Knockdown of CDC34 significantly suppresses L858R‐ and T790M-EGFR-driven lung cancer. (a) The EGFR^L858R^-transgenic mice were intranasally infected with virus particles containing shNC or sh*CDC34*, treated with DOX, and scanned by micro-CT. (b) Hematoxylin-eosin staining of the lung sections from mice of each group. (c) IHC analysis of CDC34, Ki67, and EGFR in tumors from mice of each group. (d) The expression of CDC34 and EGFR in the tumor samples from the mice of each group was detected by Western blot. (e) Kaplan–Meier survival curve of the mice. (f) The EGFR^T790M/del(exon19)^-transgenic mice were intranasally infected with viral particles containing shNC or sh*CDC34*, treated with DOX, sacrified one month later, and the lung tissue lysates were subjected to immunoblot to detect the expression of CDC34 and EGFR. (g) The EGFR^T790M/del(exon19)^-transgenic mice were intranasally infected with virus particles, treated with DOX, and scanned with micro-CT one month later. (h) Hematoxylin-eosin staining of the lung sections from mice of each group. (i) IHC analysis of CDC34, Ki67, and EGFR in lung tumors from the mice. (j) The expression of CDC34 and EGFR in the tumor samples from the mice of each group. Scale bar, 1 mm. **, *P*<0.001.

### Knockdown of CDC34 inhibits EGFR^T790M/del(E746-A750)^-driven lung cancer

The EGFR T790M mutation is associated with acquired resistance to erlotinib in NSCLCs. We tested the effects of shCDC34 on Tet-op-EGFR^T790M/Del(E746-A750)^/CCSP-rtTA transgenic mice^48^. Interestingly, we found that silencing of CDC34 (Fig. 6f) led to alleviated lung carcinogenesis in the mice, reflected by micro-CT (Fig. 6g) and histological examination (Fig. 6h). Knockdown of CDC34 also reduced Ki67 and EGFR expression (Fig. 6i, j).

## Discussion

In this study, we used the genome-wide siRNA screening to identify UPGs that are critical to lung carcinogenesis, and unveiled 31 candidates that were required for proliferation of A549 and H1975 cells (Fig. 1, Supplementary Fig. 1). Among them, CDC34 represented the most significant one, which was elevated in 67 of 102 (65.7%) NSCLCs and was inversely associated with clinical outcome of the patients (Table 1, Fig. 1). Previous studies showed that CDC34 was overexpressed in acute lymphoblastic leukemia^49^, multiple myeloma^50^, breast cancer^51^, hepatocellular carcinomas^52, 53^, and prostate cancer^54^. Therefore, the mechanisms of CDC34 in promoting the initiation and progression of malignant neoplasms warrant further investigation.

Cigarette smoking is responsible for 1.38 million lung cancer deaths each year worldwide^55^. Tobacco smoke causes genomic mutations in tumors ^56^ and counterpart normal controls^57, 58^, promotes cell proliferation, inhibits programmed cell death, and facilitates angiogenesis, invasion and metastasis potentials, and tumor promoting inflammation^59, 60^. We found that CDC34 was overexpressed in 44 of 59 (74.6%) smoker NSCLCs and in 23 of 43 (53.5%) nonsmoker patients (Table 1). In another cohort^37^, CDC34 expression in smokers was significantly higher than in nonsmokers (Fig. 1l). These results suggest that tobacco smoke may induce CDC34 expression to maintain hyperactivation of EGFR oncoprotein, contributing to lung carcinogenesis. Tobacco and tobacco smoke contain 73 carcinogens^61^, whether these compounds could modulate CDC34 expression or not remains an open question.

CDC34 functions as a K48 Ub chain-building enzyme and cognate E2 of SCF E3 ligases, and collaborates with HECT-, RING-, and RING-between-RING (RBR)-E3s to control proteolysis of substrates^62^. So far, no evidence suggests a role for CDC34 in EGFR turnover, which is tightly controlled by UbcH7/Ubc4/5/c-Cbl cascade^15, 16^. We showed that knockdown of CDC34 *in vitro* and *in vivo* resulted in downregulation of EGFR, whereas ectopic expression of CDC34 led to upregulation of EGFR and increase in its tyrosine kinase activity (Fig. 3, 6). The downstream signaling molecules were accordingly affected (Fig. 3). Mechanistically, the K11 – I86 proportion of CDC34 bound EGFR at its CR region, and competed with c-Cbl to bind EGFR and inhibited the K48-linked polyubiquitination and subsequent degradation of the substrate (Fig. 4, 5). These results demonstrate a previously unreported function of CDC34, and indicate that an E2 Ub conjugase can compete with E3 ligase to protect proteolysis of substrates.

CDC34 exerts oncogenic or tumor-promoting functions, and has been used as a therpeutic target for drug development^63^. Inhibition of CDC34 enhances anti-myeloma activity of proteasome inhibitor bortezomib^50^ and contributes to the chemopreventive activity of Chinese herbs (anti-tumor B, ATB) in murine model of lung cancer^64^. We showed that silencing of CDC34 inhibited cell proliferation and colony forming activity of NSCLC cells *in vitro* (Fig. 2, 3), and suppressed tumor growth and prolonged lifespan of xenograft NSCLC mouse models *in vivo* (Fig. 3). In an EGFR^L858R^-driven mouse lung adenocarcinoma model, sh*CDC34* treatment significantly inhibited cancer progression and prolonged survival time of the mice (Fig. 6). These results indicate that CDC34 represents a rational drug target for NSCLC. Moreover, treatment of the patients with gefitinib and erlotinib will fail because of the development of EGFR T790M mutation^65^, which affects the gatekeeper residue in the kinase catalytic domain and weakens the interaction of the inhibitors with the target^66^. Efforts have been made to overcome drug resistance^67, 68^. Here, we showed that silencing of CDC34 dramatically suppressed tumor dissemination and cell proliferation in EGFR^T790M/Del(exon 19)^-driven lung cancer (Fig. 6F). Knockdown of CDC34 downregulated the expression of EGFR in EGFR^T790M/Del(exon 19)^ mice (Fig. 6). These data demonstrate that silencing of CDC34 can overcome T790M mutation-caused drug resistance, therefore represents an attrative therapeutic target in lung cancer. Preclinical and clinical studies of using CDC34 inhibitor to treat lung cancer with or without EGFR mutation warrant intensive investigation.

## Materials and Methods

### Patient samples

The study was approved by the local research ethics committees of all participating sites; all lung cancer samples were collected with informed consent. The diagnosis of lung cancer was confirmed by at least two pathologists. Tissue samples were taken at the time of surgery and quickly frozen in liquid nitrogen. The tumor samples contained a tumor cellularity of greater than 60% and the matched control samples had no tumor content.

### siRNA library

Four Human siGENOME SMARTpool siRNA Libraries were obtained from Thermo Scientific Dharmacon (Lafayette, CO, USA). These libraries included Human Deubiquitinating Enzymes (G-004705-05), Human Ubiquitin Conjugation Subset 1 (G-005615-05), Human Ubiquitin Conjugation Subset 2 (G-005625-05), Human Ubiquitin Conjugation Subset 2 (G-005635-05), and siGENOME Controls Complete Kit (K-002800-C2-02). Transfections were performed by using DharmFECT transfection reagent (T-2001-01) according to the manufacturer’s instructions.

### Cell culture

NSCLC lines A549, H460, H1975, HCC827, and human embryonic kidney line HEK293T were obtained from the American Tissue Culture Collection (ATCC; Manassas, VA, USA) and cultured in Dulbecco modified Eagle medium (DMEM) or RPMI 1640 supplemented with 10% fetal bovine serum. Cell viability of A549 and H1975 in siRNA library screening was assayed by the CellTiter-Glo Reagent (Promega, Fitchburg, WI, USA) according to the manufacturer’s instructions. The z-score [z=(x−m)/s], where x is the raw score to be standardized, m is the mean of the plate, and s is the standard deviation of the plate, was determined for each SMARTpool within the plate ^34^. The z-scores from the three replicates for each SMARTpool were averaged and the SD determined.

### Antibodies and reagents

The antibodies used in this study were as follows: anti-β-Actin, anti-Flag (Sigma-Aldrich, St. Louis, MO, USA); anti-pEGFR (Y1173), anti-ERK, anti-pAKT and AKT (Santa Cruz Biotechnology, Santa Cruz, CA, USA); anti-pSTAT3 (Y705, S727), anti-EGFR, anti-STAT3 (Cell Signaling Technology, Beverly, MA); anti-CDC34, anti-HA (ABclonal, Cambridge, MA, USA); anti-GST (Invitrogen, Frederick, MD, USA); anti-Ki67, anti-c-Cbl (Abcam, Cambridge, UK); anti-rabbit, anti-goat or anti-mouse HRP-conjugated secondary antibody (Pierce Biotechnology, Rockford, IL, USA). Reagents used were: Chemiluminescent Western detection kit (Cell Signaling); cycloheximide (CHX) (Amresco Inc., Solon, OH, USA); MG132 and chloroquine (Sigma-Aldrich); PS-341 (Millennium Pharmaceuticals Inc., Cambridge, MA, USA).

### siRNA, shRNA, plasmids and transfections

siRNA or shRNA were purchased from GenePharmaCo.,Ltd (Shanghai, China) and the sequences are as follows: GCUCAGACCUCUUCUACGA (si*CDC34*-1#); GGACGAGGGCGAUCUAUAC (si*CDC34*-2#); GAUCGGGAGUACACAGACA (si*CDC34*-3#); UGAACGAGCCCAACACCUU (si*CDC34*-4#); GGCTCAGACCTCTTCTACGAC (human *CDC34*-shRNA-1#); GAGTGTGATCTCCCTCCTGAA (human *CDC34*-shRNA-2#); CTCTTCTACGACGACTACTAT (mouse *CDC34*-shRNA-1#); GAGTGTAATTTCGCTGCTGAA (mouse *CDC34*-shRNA-2#).

FLAG-*CDC34* vector was constructed based on pcDNA3.1 plasmid; HA-*CDC34* vector was constructed based on pCS2 plasmid. All *CDC34* mutants were subcloned from Flag or HA-tagged *CDC34* vectors. FLAG-*EGFR* was cloned from the pCAG-3Flag-HA-*EGFR* vector and FLAG-*EGFR^CR^* was subcloned from FLAG-*EGFR* vector. pFlag-CMV4-*c-Cbl* vector was kindly provided by Dr. Jianhua Mao (Shanghai Institute of Hematology, Rui Jin Hospital Affiliated to Shanghai Jiao Tong University School of Medicine, China). GST or His-tagged *CDC34*, *EGFR*, *c-Cbl* were generated based on the backbone of pGEX-4T-1 and pET28a (kindly provided by Dr. Quan Chen, Institute of Zoology, Chinese Academy of Sciences, Beijing, China), respectively. The sh*CDC34* constructs were made with PLKO.1 backbone (kindly provided by Dr. Wanzhu Jin, Institute of Zoology, Chinese Academy of Sciences) using Age I and EcoR I sites. Cells were transfected with siRNA, shRNA or plasmids using the Lipofectamine 2000 or Lipofectamine 3000 (Invitrogen).

### Lentivirus-mediated transfection

For lentiviral particle production, sh*CDC34* constructs in PLKO.1 were co-transfected with psPAX2 and pMD2G into HEK293T cells. The culture medium was replaced with fresh medium after 6 h, and the supernatants were harvested 48 h and 72 h post transfection. A549-luciferase cells were infected with viral particles in the presence of 8 μg/mL polybrene followed by puromycin selection to generate *CDC34* stably knockdown cells.

### RT-PCR

The total RNA was isolated using the TRIZOL reagent (Invitrogen) and the phenol-chloroform extraction method according to the manufacturer’s instruction. Total RNA (2 μg) was annealed with random primers at 65°C for 5 min. The cDNA was synthesized using a 1st ‐STRAND cDNA Synthesis Kit (Fermentas, Pittsburgh, PA, USA). Quantitative real-time PCR was carried out using SYBR PremixExTaq (Takara Biotechnology, Dalian, China). The primers used for quantitative RT-PCR are as follows: human *GAPDH*, 5′-GAGTCAACGGATTTGGTCGT-3′ (forward) and 5′-GACAAGCTTCCCGTTCTCAG-3′ (reverse); mouse *GAPDH* 5’-AGTATGACTCCACTCACGGCAA-3’ (forward) and 5’-TCTCGCTCCTGGAAGATGGT-3’ (reverse); human *CDC34*, 5′-GACGAGGGCGATCTATACAACT-3′ (forward) and 5′-GAGTATGGGTAGTCGATGGGG-3′ (reverse); mouse *CDC34*, 5′-CCCCAACACCTACTATGAGGG-3′ (forward) and 5′-ACATCTTGGTGAGGAACCGGA-3′ (reverse); human *EGFR*, 5′-GGACTCTGGATCCCAGAAGGTG-3′ (forward) and 5′-GCTGGCCATCACGTAGGCTT-3’ (reverse). Each sample was analyzed in triplicate for three times.

### Immunofluorescence microscopy

Cells grown on coverslip (24 mm × 24 mm) were fixed with 4% paraformaldehyde for 15 min, washed with 150 mM glycine in PBS, and permeabilized with 0.3% Triton X-100 in PBS for 20 min at room temperature. After blocking with 5% BSA, the cell smears were incubated with indicated primary antibodies overnight at 4°C, washed, and FITC/PE ‐labeled secondary antibody in PBS was added to the cell smears. Images were taken by a laser scanning confocal microscopy (Zeiss, Oberkochen, Germany).

### Immunohistochemistry analysis

IHC assay was performed with anti-CDC34, anti-Ki67 and anti-EGFR antibodies. Briefly, formalin-fixed, paraffin-embedded human or mouse lung cancer tissue specimens (5 mm) were deparaffinized through xylene and graded alcohol, and subjected to a heat-induced epitope retrieval step in citrate buffer solution. The sections were then blocked with 5% BSA for 30 min and incubated with indicated antibodies at 4°C overnight, followed by incubation with secondary antibodies for 90 min at 37°C. Detection was performed with 3, 3’-diaminobenzidine (DAB, Zhongshan Golden Bridge Biotechnology, Beijing, China) and counterstained with hematoxylin, dehydrated, cleared and mounted as in routine processing. The scoring of immunoreactivity was calculated as IRS (0–12)=RP (0–4) × SI (0–3), where RP is the percentage of staining-positive cells and SI is staining intensity.

### Western blotting

Cells were lysed on ice for 30 min in RIPA buffer (50 mM Tris-HCl pH 7.4, 150 mM NaCl, 0.1% SDS, 1% deoxycholate, 1% Triton X-100, 1 mM EDTA, 5 mM NaF, 1 mM sodium vanadate, and protease inhibitors cocktail), and protein extracts were quantitated. Proteins (20 mg) were subjected to 8–15% sodium dodecyl sulfate-polyacrylamide gel electrophoresis (SDS-PAGE), electrophoresed and transferred onto a nitrocellulose membrane. After blocking with 5% non-fat milk in Tris-buffered saline, the membrane was washed and incubated with the indicated primary and secondary antibodies and detected by Luminescent Image Analyzer LSA 4000 (GE, Fairfield, CO, USA).

### CO-immunoprecipitation

Cells were treated either with or without 10 μM MG132 for 3-4 h before lysis. After wash with cold PBS for two times, cells were suspended in IP lysis buffer (40 mM Tris-HCl, pH 7.4, 137 mM NaCl, 1.5 mM MgCl_2_, 0.2% sodium deoxycholate, 1% Nonidet P-40, 2 mM EDTA, 1 mM Na_3_VO_4_, 1 mM NaF, 1 mM PMSF, complete protease inhibitors cocktail) and cleared by centrifugation. Indicated antibody was added and incubated overnight with each cell lysate at 4°C. Protein A/G PLUS ‐Agarose beads (Santa Cruz) were added after washing for 3 times with lysis buffer. After 2-hour of incubation, beads were washed four times, 5 minutes per wash in IP wash buffer (40 mM Tris-HCl, pH 7.4, 137 mM NaCl, 1.5 mM MgCl_2_, 2 mM EDTA, 0.2% Nonidet P-40).

### Protein purification, GST or His pull-down assay, and in vitro kinase assay

GST or His-tagged CDC34, EGFR, c-Cbl proteins were expressed in *E. coli* Rosetta (DE3). GST fusion proteins were purified on glutathione-Sepharose 4 Fast Flow beads (GE Health Science, Pittsburgh, PA, USA) and His fusion proteins were purified on HisPur Cobalt Resin (Thermo Scientific, Basingstoke, UK), respectively. The GST was removed with thrombin (Amresco). For the GST pull-down, 2 μg of GST-fusion protein was incubated with cell lysates or purified proteins for 2 h at 4 °C and then washed 5 times with 1 mL PBS buffer. The precipitate complex was boiled with sample buffer containing 1% SDS for 5 min at 95 °C and subjected to SDS-PAGE. The nitrocellulose membrane was stained with Ponceau S and followed by immunoblotting with indicated antibodies. *In vitro* EGFR kinase activity was determined using Universal Tyrosine Kinase Assay Kit (TaKaRa Biotechnology), following the manufacturer’s instructions.

### Animal studies

The animal studies were approved by the Institutional Review Board of Institute of Zoology, Chinese Academy of Sciences, and the methods were carried out in accordance with the approved guidelines. For xenograft tumor model, six-week-old SCID Beige mice were maintained in the pathogen-free (SPF) conditions. A549-luciferase cells (1×10^6^) stably expressing the sh*CDC34* or shNC were injected into the lateral tail veins of the mice. After 30 days, the tumors were monitored by IVIS Spectrum Imaging System (Caliper Life Sciences, Hopkinton, MA, USA). For EGFR^L858R^ or EGFR^T790M/Del(exon19)^-driven lung tumors, 6-week-old Tet-op-EGFR^mutant^/CCSP-rtTAFVB mice (kindly provided by Professor Liang Chen, National Institute of Biological Sciences, China) were intranasally administrated with shCDC34 or shNC lentiviral particles once a day for 3 days. One week after the first lentiviral administration, DOX was added to feed the mice and the lung tumors were analysed by micro-CT (PerkinElmer, Waltham, MA, USA) scanning. Mice were anesthetized by mixture of oxygen/isoflurane inhalation and positioned with legs fully extended, and assayed according to manufacturers’ instruction. Survival of the mice was evaluated from the first day of DOX treatment until death or became moribund, at which time points the mice were sacrificed.

### Statistical analysis

All experiments were repeated at least three times and the data were presented as the mean±SD unless noted otherwise. Differences between data groups were evaluated for significance using Student’s *t*-test of unpaired data or one-way analysis of variance. *P* values less than 0.05 indicate statistical significance.

## Acknowledgements

This work was supported by the National Natural Science Funds for Distinguished Young Scholar (81425025), the National Key Research and Development Program of China (2016YFC0905500), the “Personalized Medicines——Molecular Signature-based Drug Discovery and Development”, Strategic Priority Research Program of the Chinese Academy of Sciences (XDA12010307), the National Natural Science Foundation of China (81672765), and grants from the State Key Laboratory of Membrane Biology. The study sponsor had no role in the design of the study; the data collection, analysis, or interpretation; the writing of the article; or the decision to submit for publication.

## Author Contributions

The project was conceived and designed by G.B.Z. The experiments were conducted by X.C.Z, G.Z.W., J.L., C.Z., D.L.Z., L.M., S.H.G., and L.W.Q․. Biospecimens were harvested/provided by Y.C.Z., Y.C.H., B.Z., C.L.W․. The EGFR transgenic mice were provided by L.C․. Data were analyzed by G.B.Z․. The manuscript was written by G.B.Z․.

## Disclosure of potential conflicts of interest

No potential conflicts of interest were disclosed.

